# Long-Term Reactivation of Multiple Sub-Assemblies in the Hippocampus and Prefrontal Cortex

**DOI:** 10.1101/2024.12.25.630328

**Authors:** Masami Tatsuno, Jean-Marc Fellous

**Author notes:** Corresponding Authors: Masami Tatsuno and Jean-Marc Fellous.

## Abstract

In resting or sleep periods following a task, neurons in various parts of the brain become reactivated, with firing patterns often similar to that during the task. This reactivation plays an essential role in memory consolidation. However, detecting these reactivating episodes has been challenging because not all neurons recorded may be directly relevant to the memory being consolidated. Here, we propose a novel spike train clustering (STC) method for detecting groups of neurons (clusters) with partial synchronous firing. From a mechanistic standpoint, the correlated activity of an ensemble can arise from neuronal interactions within the recorded local ensemble, common external inputs, or both. To quantify these contributions, we propose to use information geometry (IG) by taking advantage of its capacity for orthogonal decomposition of neural interactions. We analyzed simultaneous single unit activity from rat medial prefrontal cortex (mPFC) and area CA1 of the hippocampus when animals explored novel objects and when they slept before and after the exploration. We demonstrate that multiple reactivations by different subsets of neurons (clusters) occurred. Those reactivations could extend over 11 hours, the entire recording duration of post-task rest after the exploration epoch. Long-lasting reactivation was not detected when all neurons were included as a single cluster in the analysis. We also showed that pairwise interactions in reactivating clusters tended to be more strongly modified than in non-reactivating ones. In addition, the pairwise interactions of the reactivating clusters in CA1 were strongly modulated by the task experience but not in the mPFC. These results indicate that hippocampal reactivation following novel experience is likely to be induced within the hippocampal circuits. In contrast, mPFC reactivation is likely to be driven by external inputs, possibly in part from the hippocampus.

## 1 Introduction

Current techniques allow for the simultaneous recording of hundreds to thousands of neurons from different brain areas in behaving animals (Buzsaki, 2004; Miller and Wilson, 2008; Nicolelis and Lebedev, 2009; Rector et al., 2009; Ahrens and Engert, 2015; Buzsaki et al., 2015; Giocomo, 2015; Grinvald and Petersen, 2015; Hamel et al., 2015; Tatsuno, 2015; Harris et al., 2016; Kim et al., 2016; Steinmetz et al., 2018; Giri et al., 2019; Stringer et al., 2019; Urai et al., 2022; Stringer and Pachitariu, 2024; Yuste et al., 2024). It has been conjectured that information processing relies on the dynamic formation of neural assemblies (Hebb, 1949), which, in practice, remain challenging to identify and study experimentally as well as theoretically (Russo and Durstewitz, 2017; Mackevicius et al., 2019; Watanabe et al., 2019; Yuste et al., 2024). The quantification and interpretation of the possible sources of correlation associated with these assemblies have also been a difficult problem (de la Rocha et al., 2007; Nienborg and Cumming, 2010; Pospisil and Bair, 2021) because correlation can arise from local synaptic interactions within the ensemble of recorded neurons or from correlated external inputs to the neurons, or a mixture of both. The development of a systematic detection algorithm of neuronal clusters and the quantification of the possible sources of their correlations is, therefore, a critical step towards the analysis and interpretation of large-scale multi-neuronal spike activity. Most studies of reactivation match the pairwise firing correlation patterns occurring during the task to that occurring immediately after post-task sleep onset and find relatively short periods of reactivation (less than 30 minutes). A seminal study however used ICA applied on the full population of recorded CA1 neurons to provide evidence for long-term reactivation in the hippocampus (Giri et al., 2019). The tasks used in this study were on linear tracks, providing a restricted spatial extent that might have increased pairwise correlation and hence reactivation, cell assemblies were defined on the bases of firing rate variance (ICA) rather than correlation, and no comparison was made across different brain areas. Adding to and complementing this body or work, we propose a novel analysis framework by combining spike train clustering (STC) (Fellous et al., 2004) and information geometry (IG) (Amari and Nagaoka, 2000). STC identifies neuronal subgroups that exhibit partial synchrony using fuzzy-clustering methods. STC is based on spike timing (Humphries, 2011; Lyttle and Fellous, 2011) rather than spiking order(Watanabe et al., 2019). IG estimates the pairwise correlation strengths independently from the amount of external inputs the assembly receives (Tatsuno and Okada, 2004; Tatsuno et al., 2009; Nie and Tatsuno, 2012). IG has also been shown to be robust to oscillatory brain states (Nie et al., 2014b), and its properties in up to 10 neuron interactions have been investigated analytically (Nie et al., 2014b). By applying the proposed method to long-term recordings from rat medial prefrontal cortex (mPFC) and area CA1 of the hippocampus, we investigate the following specific questions: Do all or subsets of recorded neurons reactivate after an object-novelty task? Are reactivation dynamics different for different subsets? Do strongly reactivating clusters involve more task-related neuron pairs? Are reactivations in mPFC and CA1 primarily driven by synaptic changes or external inputs? Some of the results of this study were presented in abstract form elsewhere (Lipa et al., 2006; Lipa et al., 2007).

## 2 Results

### 2.1 Spike train clustering identifies a subset of neurons with partial synchrony

Spike train clustering (STC) was initially proposed to analyze the response of a single neuron to repeated presentations of the same stimulus (Fellous et al., 2004). It was shown to be effective at identifying subgroups of trials that exhibited common spike patterns(Humphries, 2011). In principle, this method is applicable to any temporally time-locked ensemble of spike trains that were recorded from one neuron or an ensemble of neurons. To investigate the applicability of this approach to the multi-neuronal case, we built a biophysical simulation of ten cortical principal neurons using the NEURON simulator (Hines and Carnevale, 1997; Tanaka, 2000), where five neurons were weakly interconnected, forming a cluster (Fig. 1A). Representative traces of the membrane voltage of a cell that belongs to the cluster and one that does not are shown (Fig. 1A). These traces are indistinguishable by commonly used statistics of the activity of single neurons, such as the mean, standard deviation, coefficient of variation and Fano factor of the membrane potentials or inter-spike intervals. A three-second spike raster of spontaneous activity of all ten neurons also shows that no obvious synchrony can be seen between neurons within the cluster (gray circles) and those outside (open circles) (Fig. 1B). The STC algorithm was then applied to this spike raster and two groups of spike trains were identified (Fig. 1C). The re-ordered spike raster shows that the two clusters correspond exactly to the interconnected group of neurons (below the red dashed line) and the independent group of neurons (above the red dashed line). Furthermore, partially synchronous spike firing in the connected group becomes visually obvious (ovals). In summary, we demonstrate that the STC is applicable to multi-neuronal activity and that it identifies subsets of neurons with partial synchrony.

**Figure 1:**
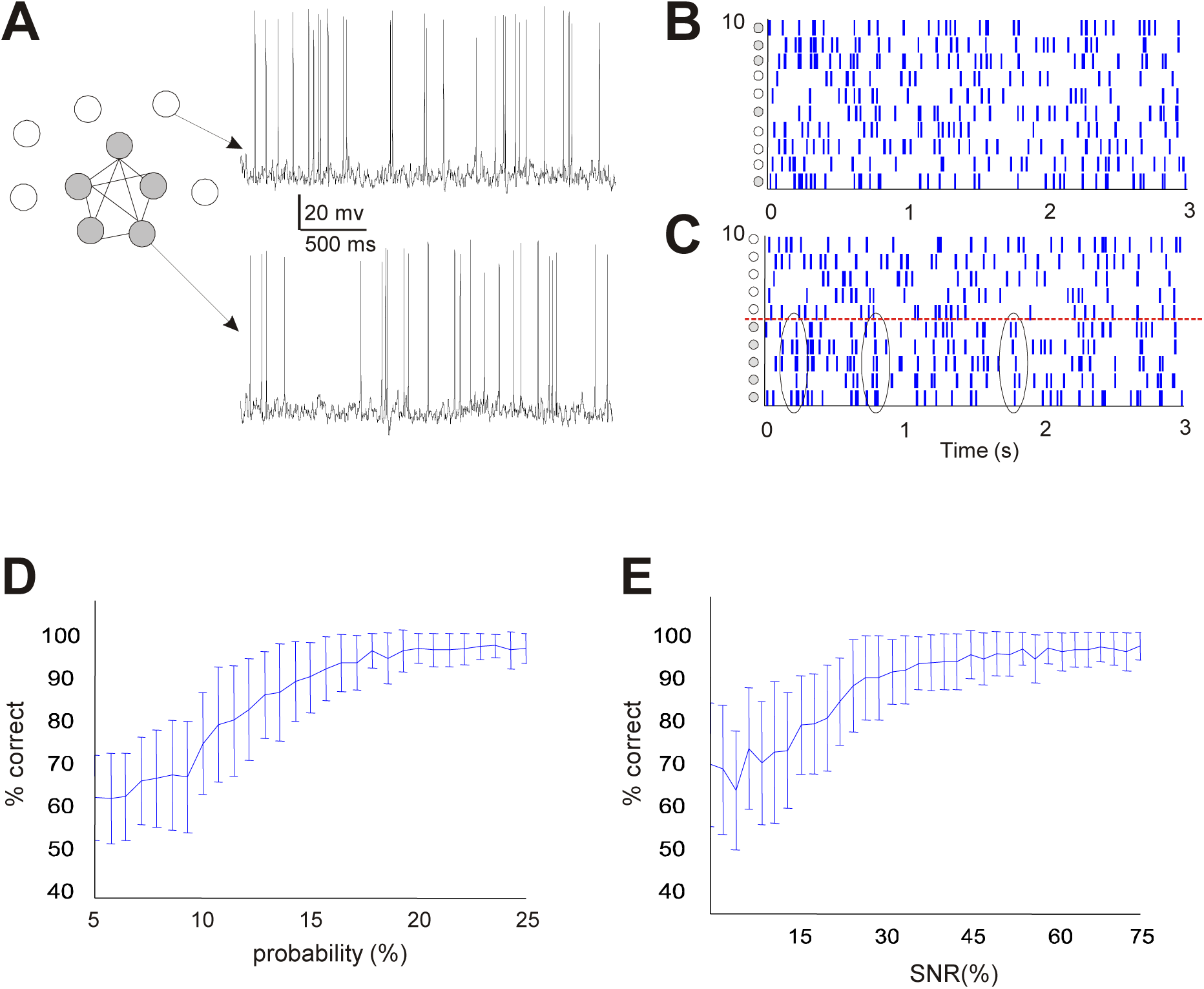
Biophysical simulation and performance of STC. **(A)** Architecture of the network. Five cells are fully interconnected (grey), and five others fire independently (open). The membrane potential is shown for a cell outside the assembly (top) and for a cell within the assembly (bottom). **(B)** Rastergram of spontaneous activity for 3 seconds. **(C)** Same rastergram as in **(B)**, but cells are clustered by STC into two groups (horizontal dashed line). The symbols to the left of the Y axis in **(B)** and **(C)** represent the neurons in **(A)**. **(D)** Performance of STC with modulation of the average connection strength, expressed as the probability for a presynaptic spike to elicit a spike postsynaptically. **(E)** Performance of STC with modulation of common inputs. In **(D)** and **(E)**, error bars are standard deviations of the percent correct classification obtained from 50 independent simulations.

### 2.2 Spike train clustering efficiently detects subsets of neurons formed by synaptic interactions or correlated inputs

To investigate the performance of STC further, we examined whether STC could detect partial synchrony due to synaptic interconnections as well as common inputs from an external source. In the first case, the detection performance was established based on 3 seconds of data for cells that fired at about 7 Hz (coefficient of variation about 0.7). Here, we express connection strength as the probability that a presynaptic cell fires its postsynaptic target with a single action potential. For simplicity, connections between cells of the cluster are implemented as weak AMPA synaptic conductance. Since cells received uncorrelated background noise, the probability for one presynaptic cell to fire its target depended not only on their direct synaptic connection but also on the background noise and the indirect multi-synaptic pathways involving the other cells within the cluster. The notion of connection strength used here should, therefore, be regarded as a statistical feature of a pair of recorded neurons rather than an anatomical statement regarding the presence of a direct monosynaptic connection. The connection strength is measured empirically as the number of postsynaptic spikes emitted in response to 100 single spikes from the presynaptic cell (expressed as a percentage). When the connection strength is systematically varied in the network, the detection of the cluster was significantly above chance for connection strength values that were within that observed in vivo (10% in Fig. 1D) (Bartho et al., 2004; Song et al., 2005).

In the second case, partial synchrony was due to common inputs. The neurons were interconnected by very weak synapses (< 2% firing probability of a postsynaptic neuron due to a single presynaptic spike) but received part of their average background inputs from a common source. The common input was simulated as a modulation of the background excitatory synaptic noise, expressed as a percentage of the noise level. Using the same method as above, 15% modulation of the background inputs could be effectively detected above chance (Fig. 1E). Note that in these simulations, the overall firing rate of all cells in the assembly was kept constant and indistinguishable from that of the cells outside the cluster, so as not to bias the algorithm into clustering cells by firing rates rather than by partial synchrony. In summary, our investigation using a network of biophysical model neurons showed that STC could efficiently detect neuronal clusters that are formed by weak synaptic interactions and by weak common correlated inputs.

### 2.3 Information geometric measures can estimate the effect of synaptic interactions and external inputs separately

Information Geometry (IG) has emerged from studies of geometrical structures involved in statistical inference (Amari and Nagaoka, 2000; Amari, 2016). Among IG’s applications to various fields of information sciences, it has been shown that IG provides a useful tool for the analysis of correlations in multi-neuronal spike data (Amari, 2001; Nakahara and Amari, 2002; Tatsuno et al., 2009; Nie and Tatsuno, 2012; Shimazaki et al., 2012; Nie et al., 2014b, a). By representing neuron’s activity as binary variables, the second-order information geometric measure (IGM) between neurons *i* and *j*, denoted by *θ*_*ij*_, was shown to be statistically independent from the modulation of the mean firing rate of each neuron (Amari, 2001; Nakahara and Amari, 2002; Amari, 2009). This is one of the key advantages of the IG approach over traditional measures, such as the Pearson correlation coefficient. In addition, the first-order IGM of neuron *i*, denoted by *θ*_*i*_, is linearly related to the amplitude of external inputs to the neuron (Tatsuno et al., 2009; Nie and Tatsuno, 2012).

To validate the IG approach in biologically plausible conditions, we calculated *θ*_*ij*_ and *θ*_*i*_ from spike trains generated by biophysical simulations of ten cortical neurons, which were used to assess the STC performance. We examined whether IG could distinguish pairwise neural interactions due to the synaptic connections or external inputs. We considered two scenarios. In the first scenario, the AMPA synaptic conductance was increased discretely, but external inputs remained the same. This corresponds to a simulated Hebbian learning case. In the second scenario, the amplitude of external inputs was increased discretely, but the AMPA synaptic conductance remained the same. This corresponds to a non-learning case. For clarity, we divided the simulations into three epochs of three-second duration, between which the synaptic connections or the external inputs were changed in a discrete manner.

In the first learning scenario, the mean firing rate of the five connected neurons increased discretely, in accordance with the discrete increase of synaptic conductance (Fig. 2A, top). The correlation coefficient (CC), a commonly used pairwise interaction measure, also increased in a discrete manner, suggesting that the CC tracked the change of synaptic conductance (Fig. 2A, middle). We then computed the information geometric measures, *θ*_*ij*_ and *θ*_*i*_. We found that *θ*_*ij*_ increased in a discrete manner (Fig. 2A, bottom, blue trace) whereas *θ*_*i*_ remained almost constant (Fig. 2A, bottom, red trace). Because the AMPA synaptic conductance was increased discretely, but external inputs remained the same, this result demonstrates that *θ*_*ij*_ and *θ*_*i*_ were able to estimate the change of synaptic conductance and the change in amplitude of external inputs separately.

**Figure 2:**
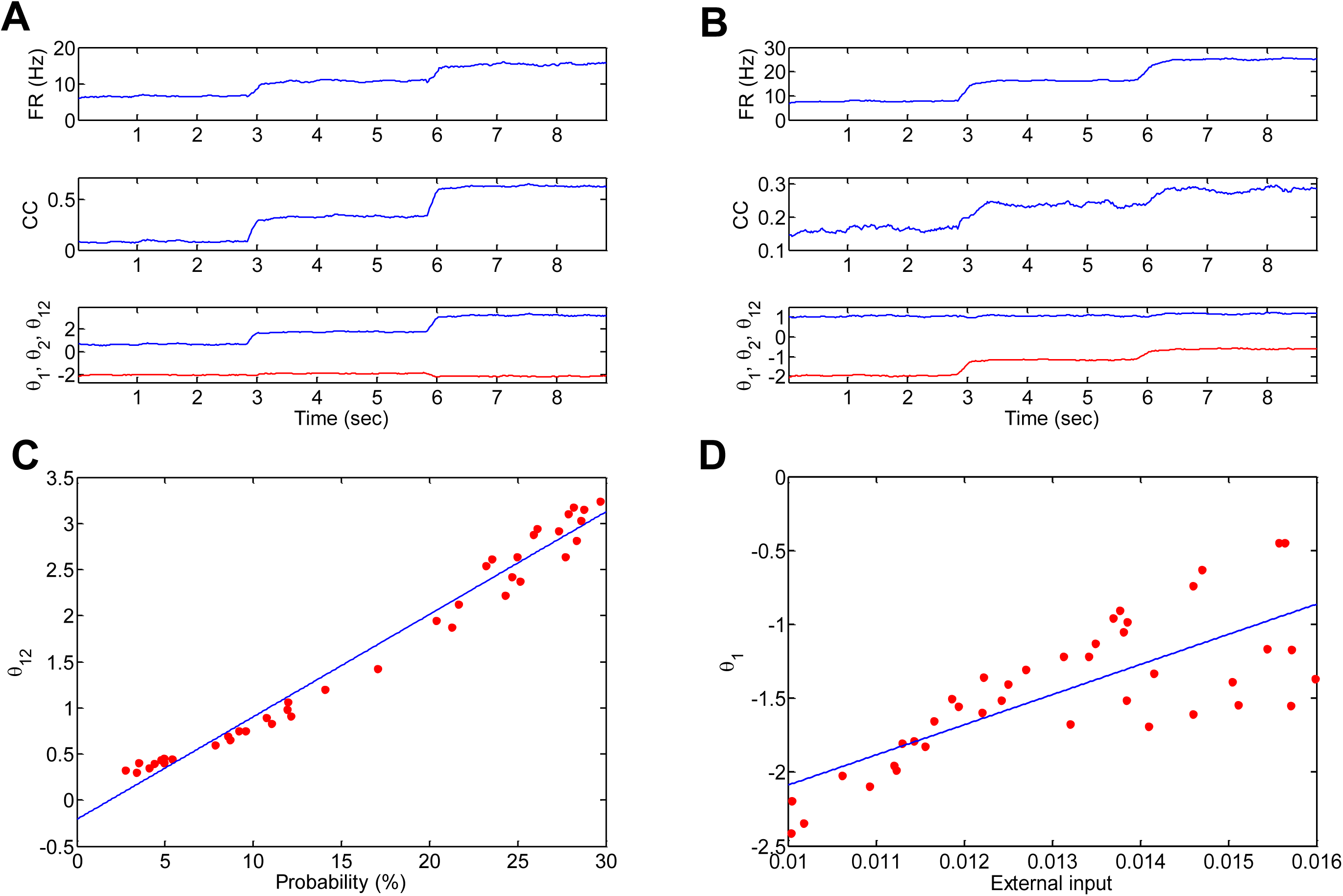
Estimation of connection strengths and external inputs by IG. **(A)** Connection strengths between the neurons in the cluster (Figure 1 **(A)**) are increased in three discrete steps. The top, middle, and bottom panels show the average firing rate, the correlation coefficient (CC), and *θ*_*ij*_ (blue) and *θ*_*i*_ (red), respectively. Note that *θ*_*i*_ and *θ*_*j*_ are identical in this analysis. 500 independent 9 seconds long simulations (each step is 3 seconds) are used for the estimation of the probability distribution. **(B)** External inputs are increased in three discrete steps. Each panel is the same with **(A)**. **(C)** Relationship between the connection strengths and *θ*_*ij*_ when connection strengths and external inputs are modulated simultaneously (the coefficient of determination = 0.98). **(D)** Relationship between the external inputs and *θ*_*i*_ for the same condition as in **(C)** (the coefficient of determination = 0.58). In **(C)** and **(D),** 40 datasets are independently generated. Each dataset consists of 500 3-second-long simulations.

In the second non-learning scenario, we observed that the mean firing rate of the five connected neurons increased discretely, tracking the discrete increase of the external inputs (Fig. 2B, top). Note the similar time course of the firing rates in both scenarios (Fig. 2A top and Fig. 2B top). This suggests that the firing rates are influenced by both the connection weights and external inputs. The CC also increased in a quasi-discrete manner as the external inputs were increased (Fig. 2B, middle). The results in the two scenarios demonstrate that the CC was affected similarly by both the synaptic connections and the firing rates. That is, the CC cannot clearly distinguish the underlying causes, as was suggested by previous theoretical investigations (Amari, 2001; Nakahara and Amari, 2002; Tatsuno and Okada, 2004). Unlike the CC, we found that *θ*_*ij*_ remained constant (Fig. 2B, bottom, blue trace) and that *θ*_*i*_ increased discretely (Fig. 2B, bottom, red trace). Taken together, these results show that *θ*_*ij*_ and *θ*_*i*_ can estimate the changes of synaptic conductance and external inputs, respectively, under conditions of biologically plausible simulations. This property is a significant advantage of the IGMs for neural data analysis.

### 2.4 Information geometric measures provide robust estimations of the strength of synaptic interactions and external inputs

In the previous section, the property of the IGMs was investigated by changing the synaptic conductance (scenario 1) and the external inputs (scenario 2) separately. Here we investigate the robustness of the IGM estimation when the strengths of the synaptic interactions and those of the external inputs were modified simultaneously.

The value of the unitary synaptic conductance and the average amplitude of the external inputs were randomly selected, and a three-second epoch of neural firing was generated. By repeating the procedure, 40 epochs of firing data with different synaptic conductance and external input amplitude were produced. Then, *θ*_*ij*_ and *θ*_*i*_ were calculated for each epoch. We found that *θ*_*ij*_ was almost linearly related to the synaptic conductance, measured by the number of postsynaptic spikes emitted in response to 100 single spikes from the presynaptic cell (Fig. 2C). We also found that *θ*_*i*_ exhibited a quasi-linear relationship with the average amplitude of the external inputs (Fig. 2D).

Unlike the CC that is bound between -1 and 1, the IGMs can take various values (e.g., 0 < *θ*_*ij*_ < 3.5 in Fig. 2C and −2.5 < *θ*_*i*_ < 0 in Fig. 2D). It is not easy to interpret those values because they are influenced by various biophysical factors in a complex manner. However, the result in this section demonstrates that a relative change of synaptic conductance and external inputs can be separately estimated by the IGMs, even when they are modulated simultaneously.

### 2.5 Short-lasting reactivation of all recorded pyramidal neurons was detected by the explained variance analysis

We applied the combined STC and IG technique to multi-unit recording data from behaving rats that explored novel objects (Ribeiro et al., 2004; Tatsuno et al., 2006). Briefly, we analyzed datasets of continuous 25-hour recordings obtained from five rats. Multi-unit spike data were recorded from the medial prefrontal cortex (mPFC) in two rats (rats 1 and 2), from area CA1 of the hippocampus in two rats (rats 3 and 4), and from the mPFC and the CA1 simultaneously in one rat (rat 5). The recording sessions consisted of a pre-task epoch (12 hours), a task epoch when the rat explored novel objects (1 hour), and a post-task epoch (12 hours). A subset of the data from four rats (rats 1, 2, 3 and 5) was used in previous publications (Tatsuno et al., 2006; Schwindel et al., 2014).

The strength of memory reactivation was estimated by the explained variance (EV) method (Kudrimoti et al., 1999). EV measures the similarity of correlation matrices between the task and post-task epochs, factoring out pre-existing correlations from the pre-task epoch. A series of 15-minute-blocks in the pre- and post-task epochs as well as the actively exploring periods of the task epoch were used to calculate EV values. Significance of EV was assessed using reversed EV by setting the threshold as (mean + 2 standard deviation) (Pennartz et al., 2004; Tatsuno et al., 2006). As the present study focuses on sleep reactivation, only the 15-minute-blocks containing at least 2/3 of post-task sleep periods (sleep-dominated blocks) were included in the significance estimation.

Using all recorded pyramidal neurons, we found that the majority of mPFC reactivation was detected up to about three hours after the onset of the post-task epoch (Fig. 3A, left, red trace). The significant reactivation in sleep-dominant blocks (gray background in Fig. 3) is indicated by asterisks (*) on the horizontal time axis. Although sporadic mPFC reactivation was observed at later times in two recordings (rats 1 and 2), strong reactivation was confined mostly within the first three hours. A similar short-lasting reactivation was obtained for the CA1 data sets (Fig. 3B, left, red trace). Most reactivation was observed typically within the first 30 minutes of the post-task epoch, which is consistent with previous study (Kudrimoti et al., 1999; Tatsuno et al., 2006) but also see (Giri et al., 2019).

**Figure 3:**
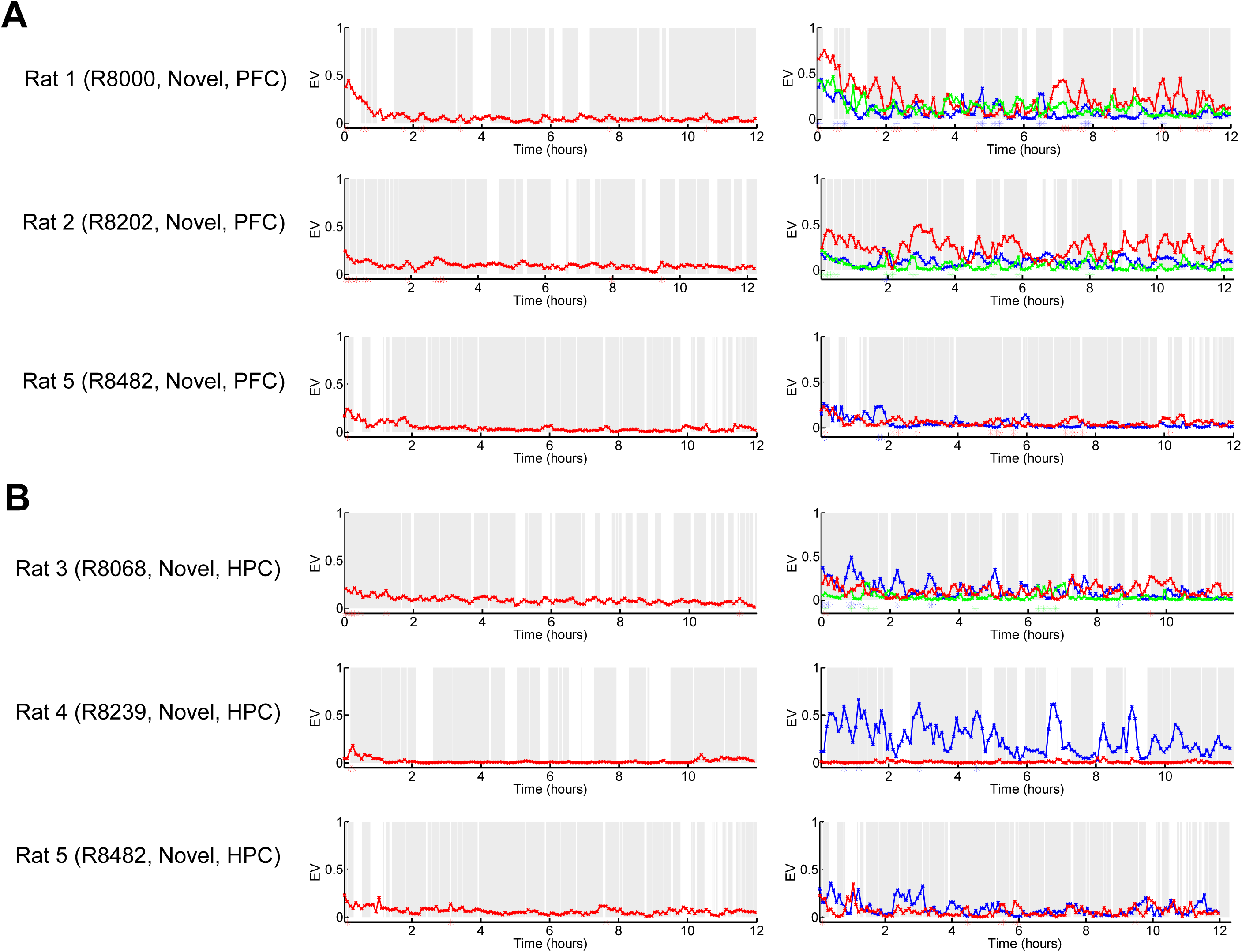
Memory reactivation analysis with and without clustering information. **(A)** (left) EV analysis of mPFC neurons without clustering. (right) EV analysis of mPFC neurons with clustering information. EV was computed by shifting a 15-minute block of post-task and pre-task correlation matrices with 5-minute steps. The significance of EV was assessed using reversed EV (mean + 2 standard deviations). **(B)** (left) EV analysis of CA1 neurons without clustering. (right) EV analysis of CA1 neurons with clustering information. The gray background areas indicates quiet periods. Star symbols on the x-axes show the time points of significant EV values.

In summary, the EV analysis using all recorded pyramidal neurons confirmed that strong reactivation could be detected for a relatively short duration, typically up to about three hours in mPFC and about 30 minutes in CA1.

### 2.6 Long-lasting reactivation of STC-clustered pyramidal neurons is detected by the explained variance analysis

We next examined if subgroups (clusters) of recorded neurons reactivate with a different time course. We applied STC to neuronal activity during the task epoch and clustered it into subgroups. The optimal number of clusters was determined by computing the ratio between the inter-cluster distance and intra-cluster distance (Fellous et al., 2004). Selecting the clusters with the largest ratio yielded two or three subgroups of the recorded pyramidal neurons. Table 1 summarizes the number of neurons and the number of neuronal pairs in the original dataset and each cluster.

**Table 1:**
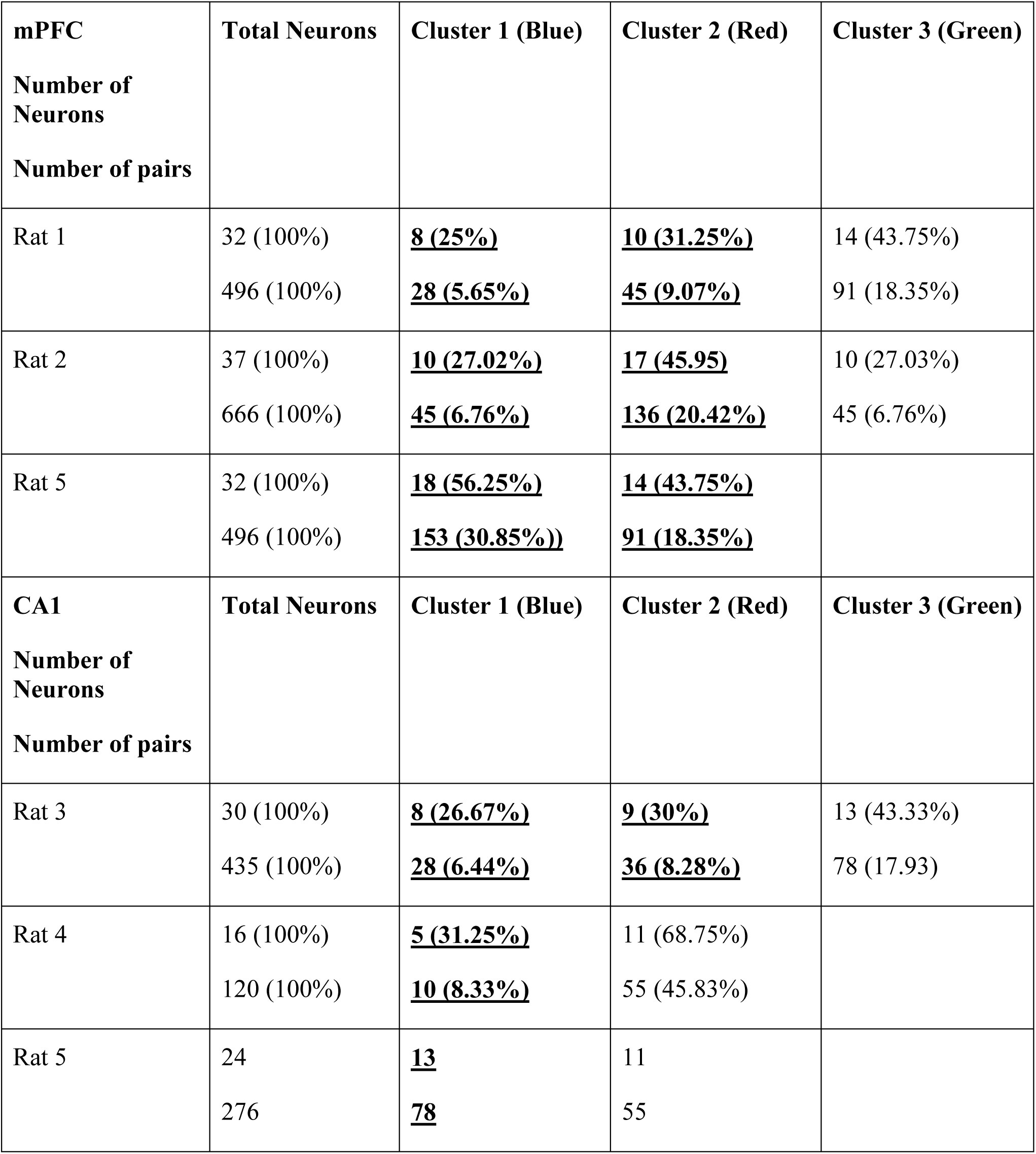
The number of neurons and the number of neuronal pairs in the original dataset and each cluster. The top half is for three mPFC rats, and the bottom half is for three CA1 rats. The top and bottom rows in each element represent the number of neurons and the number of neuronal pairs, respectively. Percent in parenthesis is in reference to the total number of neurons and the total number of neuronal pairs in each rat. Bold and underlined values indicate reactivating clusters.

The reactivation of each cluster was estimated by the EV method. A cluster that exhibits at least two consecutive reactivations or three single reactivations at the minimum was considered a reactivating cluster. We found that different clusters exhibited different time courses of reactivation (Fig. 3A, right column). In the mPFC, several clusters (blue and red in rats 1, 2, and 3) reactivated in the initial period of the post-task epoch and continued to reactivate until later in the recording, up to about 11 hours. In CA1, several clusters (blue and red in rat 3, and blue in rats 4 and 5) also exhibited long-lasting reactivations with different time courses (Fig. 3B, right column).

Long-term reactivation was not apparent when all pyramidal neurons were included in the analysis (Figs. 3A and 3B, left column). This is possibly because the EV was weakened by the combinatorically dominant pairwise correlations that were not directly related to behavior. In summary, our analysis demonstrated that the STC improved the detection of significant reactivation epochs per cluster and that those epochs could continue to occur for many hours. These rich reactivation dynamics, both in the mPFC and the CA1, are suggestive of complex underlying network interactions that selectively involve different groups of neurons (Prut et al., 1998; Chang et al., 2000).

### 2.7 Task-modulated pairwise interactions were sustained for many hours in the CA1 but not in the mPFC

To investigate how pairwise neural interactions were modified in the clusters, we computed *θ*_*ij*_for every neuron pair within each cluster, for all 15-minute blocks, as in the EV analyses. The neuron pairs were then sorted according to the strength of modulation of *θ*_*ij*_before and after the task; we first calculated *θ*_*ij*_ values in the 15-minute sleep-dominated blocks within one-hour-epochs immediately before and after the task. For each neuron pair, this procedure generated two distributions of *θ*_*ij*_ values, each corresponding to pre- and post-task one-hour epochs. We then evaluated the strength of the modulation of *θ*_*ij*_ by statistical significance using the Wilcoxon signed rank test. The neuron pairs were sorted into three groups: group 1 with the significantly positive change from the pre-task to the post-task epochs, group 2 with the non-significant change, and group 3 with the significantly negative change from the pre-task to the post-task epochs. An alpha value was set at 0.05.

To examine how the task-modulated correlations were sustained in the mPFC and CA1, the pre- and post-task sleep epochs were divided into one-hour intervals. We took the *θ*_*ij*_-distribution in the hour immediately before the task as control. The significance of *θ*_*ij*_-distributions in other hour intervals were estimated by the Wilcoxon signed rank test. To distinguish the positive and negative differences, the p-values of this test were transformed into significance scores: (1-p) for positive differences and - (1-p) for negative differences. That is, the positive and negative significances corresponded to (1-p) > 0.95 and –(1-p) < -0.95, respectively. The results were summarized as matrices of the *θ*_*ij*_ significance- scores for each cluster (Figs. 4A and 4B). Rows correspond to neuron pairs, and columns correspond to one-hour blocks. Elements in the matrix are the significance score in the range of [-1,1], color-coded from negative significance (blue) to positive significance (red). The one-hour epoch immediately before the task appears as a green column (significance score of 0) because it serves as the control epoch for the analysis. The task epoch was excluded from the matrix. The rows are sorted from group 1 to group 3, in decreasing order of the significance score between the one-hour-epochs immediately before and after the task.

**Figure 4:**
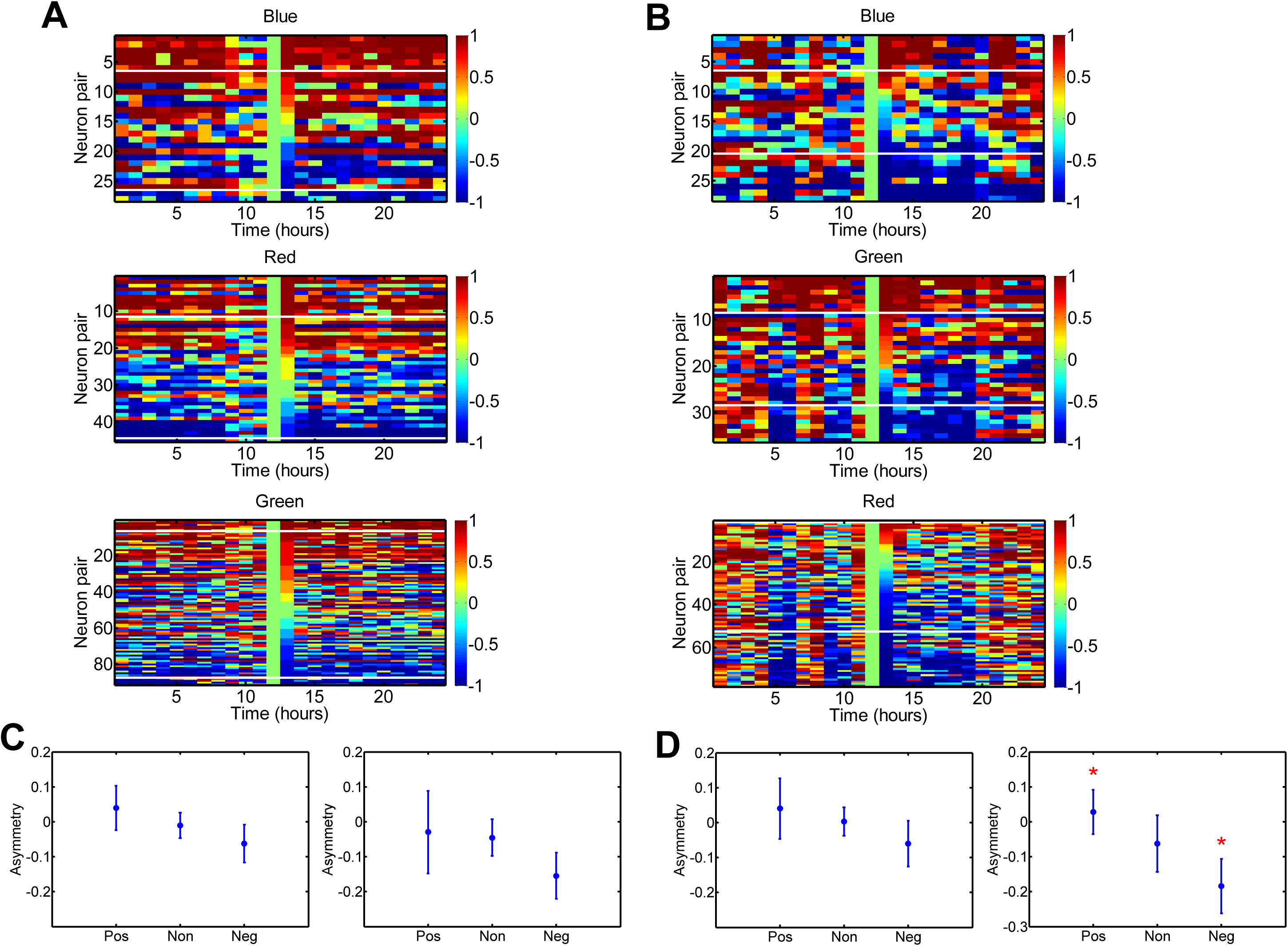
Task-dependent change of neural interactions in mPFC and CA1. **(A)** Representative mPFC recording (Rat 1) is shown. Using IG, pairwise neural interactions were estimated for all 15-minute blocks (5-minute steps). A significant change in each 1-hr block was estimated against the one-hour interval just before the task. Blue, red, and green correspond to the three clusters in Fig. 3(A) (right column, top), respectively. White horizontal lines separate the three groups (positive, non-significant, and negative changes). **(B)** Representative CA1 recording (Rat 3) is shown. Blue, red, and green correspond to the three clusters in Fig. 3(B) (right column, top), respectively. **(C)** Distribution of the asymmetry measure of the three groups (positive, non-significant, and negative changes) was plotted as mean ± sem for eight mPFC clusters (left) and for seven CA1 clusters (right). **(D)** Distribution of the asymmetry measure of the three groups (positive, non-significant, and negative changes) was plotted as mean ± sem for six reactivating mPFC clusters (left) and for four reactivating CA1 clusters (right). Stars represent significant differences with p < 0.05.

In an illustrative mPFC recording (Fig. 4A, rat 1), the ratio of significantly modulated pairwise neural interactions (group 1 + group 3) was 28.6% and 26.7% for the two reactivating clusters (blue and red traces in Fig. 3A, top, right), respectively. The corresponding ratio for the non-reactivation cluster (green trace in Fig. 3A, top, right) was 11.0%. In an illustrative recording from CA1 (Fig. 4B, rat 3), the ratio of significantly modulated pairwise neural interactions was 50.0% and 44.4% for the two reactivating clusters (blue and red traces in Fig. 3B, top, right) while that for the non-reactivating cluster (green trace in Fig. 3B, top, right) was 34.6%. These results suggest that the reactivating clusters recruit more neuron pairs that are modulated significantly. Thus, we calculated the ratio of significantly modulated pairwise interactions for all datasets (Table 2). For the animals that have both reactivating and non-reactivating clusters (mPFC: rats 1 and 2, CA1: rats 3, 4, and 5), we found that reactivating clusters recruited more neuron pairs that were modulated significantly in four out of five cases (mPFC: rats 1 and 2, CA1: rats 3 and 4). However, no significant difference was found between reactivating and non-reactivating clusters (Ranksum test; p = 0.835 for all datasets; p = 0.262 for datasets with three clusters, p = 0.533 for datasets with two clusters; p = 0.643 for mPFC datasets; p = 0.400 for CA1 datasets). This is because there was a large variance in the ratio, from 7.8% to 70.0% for reactivating clusters and from 11.0% to 38.2% for non-reactivating clusters.

**Table 2:** The ratio of significantly modulated pairwise neural interactions in each cluster. The strength of pairwise neural interactions was estimated by IG. The top half is for three mPFC rats, and the bottom half is for three CA1 rats. Bold and underlined values indicate reactivating clusters.

| **mPFC** | **Cluster 1 (Blue)** | **Cluster 2 (Red)** | **Cluster 3 (Green)** |
| --- | --- | --- | --- |
| Rat 1 | <u>**28.6%**</u> | <u>**26.7%**</u> | 11.0% |
| Rat 2 | <u>**31.1%**</u> | <u>**29.4%**</u> | 22.2% |
| Rat 5 | <u>**7.8%**</u> | <u>**8.8%**</u> |  |
| **CA1** | **Cluster 1 (Blue)** | **Cluster 2 (Red)** | **Cluster 3 (Green)** |
| Rat 3 | <u>**50.0%**</u> | <u>**44.4%**</u> | 34.6% |
| Rat 4 | <u>**70.0%**</u> | 38.2% |  |
| Rat 5 | <u>**10.3%**</u> | 11.0% |  |

Another interesting observation of the illustrative recordings was that the pairwise neural interactions appeared more strongly modulated by the task experience in CA1 than the mPFC. For example, in the top panels in Figs. 4A and 4B, the neuron pairs above the first white horizontal line are the ones that exhibit a significant positive change from the 1-hour epoch immediately before the task to the 1-hour epoch immediately after the task. For the mPFC (Fig. 4A), many of these elements are also red before the task, suggesting that they tended to exhibit positive interactions even without the task experience. For CA1 (Fig. 4B), there is the fewer number of red elements before the task experience. Then, more red elements appeared after the task experience and were sustained for a couple of hours.

To quantify the changes across the task, we computed the asymmetry index (AI) of the *θ*_*ij*_ significance- score matrices between the pre-task and post-task epochs. The AI was defined as the difference between the sum of the significance scores of all the pre-task epochs and that of all the post-task epochs (Post - Pre), normalized by the total number of the matrix elements,

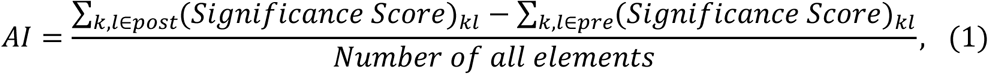

where 𝑘 runs over neuron pairs and 𝑙 runs over one-hour epochs. The one-hour epochs immediately before and after the task (the epochs used for sorting neuron pairs) were excluded from the calculations. The positive (negative) AI means that, on average, the pairwise neural interactions became more positive (negative) after the task experience.

The AI was computed for the three groups separately (group 1 with significantly positive change, group 2 without significant change, and group 3 with significantly negative change). There are eight mPFC clusters (eight traces in Fig. 3A, right panel) and seven CA1 clusters (seven traces in Fig. 3B, right panel). That is, for each of the three groups, there are eight AI values for the mPFC and seven AI values for CA1. We calculated the mean and the standard error for each group and plotted them in Fig. 4C. The negative slope from groups 1 to 3 indicates that the modulation of pairwise interactions was consistent with the group identification using the one-hour-epochs immediately before and after the task. However, the trend was not significant either in the mPFC or CA1 (Friedman’s test; p = 0.135 for the mPFC; p = 0.276 for the CA1). Next, we focused on the reactivating clusters only. The analysis included six mPFC clusters and four CA1 clusters. The statistical test showed that the mPFC was not significant (Friedman’s test, p = 0.311), but CA1 was (Friedman’s test, p = 0.038) (Fig. 4D). The post-hoc multiple comparisons showed that groups 1 and 3 were significantly different (multiple comparisons with Tukey’s honest significant difference criterion, p = 0.035, star symbol in Fig. 4D). In summary, these results suggest that the pairwise neural interactions of the reactivating clusters were strongly modulated by the task experience in CA1 but not in the mPFC.

## 3 Discussion

Reactivation of previous experiences during sleep plays a crucial role in memory consolidation. Detection of reactivating signals has been challenging, however, because only subsets of recorded neurons may be directly involved in task experiences. Estimation of neural interactions has also been difficult because pure interactions are not easily separable. Here, we proposed a novel analysis approach by combining STC and IG, where STC detects groups of neurons (clusters) with partial synchronous firing, and IG estimates neural interactions independently from the firing rates.

Application of STC to single-unit recordings of the rat mPFC and CA1 demonstrated that spiking activities in both areas were clustered on the basis of their partial pairwise correlation during a task. Only the subset of clusters reactivated after the task, and the reactivating clusters had different long-lasting dynamics up to about 11 hours after the task. Investigations of neuronal interactions by IG indicated that the reactivating clusters tended to recruit more tasked-modulated neuron pairs than non-reactivating clusters, though the effect was not statistically significant. The pairwise neural interactions were more strongly modulated by the task in CA1 than the mPFC, suggesting that CA1 reactivation could be driven by intrinsic changes within the hippocampal circuits, but mPFC reactivation could be driven by external inputs, possibly in part from the hippocampus.

Our novel object recognition study is consistent with the standard memory consolidation hypothesis, which states that novel memory is initially encoded in the hippocampus and is gradually transferred to the neocortex (McClelland et al., 1995; Buzsaki, 1998). In reality, the interaction between the hippocampus and neocortex can be more complex depending on factors such as the age of memory and the stage of sleep. It is interesting to investigate how single-unit activity for familiar experience can be clustered in the neocortex and hippocampus and how their reactivation dynamics and neural interaction change over long hours. Reactivation of hippocampus-dependent memory has been predominantly reported during slow-wave sleep, but REM reactivation should be revisited with more advanced analysis approaches.

The study showed that the dialog between the hippocampus and neocortex continues far longer than is usually assumed. The mechanism for intermittent long-lasting reactivation is not well understood. However, it may be induced by multiple waves of Arc protein expression at later times after the completion of a task (Ramirez-Amaya et al., 2005). In a previous study, Giri et al. reported that hippocampal reactivation extends for several hours after a novel experience (Giri et al., 2019). They assessed the strength of reactivation using the same EV approach. One difference between the two studies is that Giri et al. detected single long-lasting reactivation using all recorded neurons, whereas our study detected different reactivation dynamics for different neuronal clusters. With all recording neurons, our analysis showed relatively short-lasting reactivation (Fig. 3B, left). The difference between the two studies could come from various factors, such as the different implementation of novel experiences and the differences in the recording systems, spike sorting, and rat species. Overall, both studies detected long-lasting reactivation after novel experiences, and they support the importance of reactivation during sleep for memory consolidation.

It has been known that pyramidal cells in CA1 have almost no direct connections, but they receive strong inputs from CA3. The change of neuronal interaction in CA1 could reflect changes induced in CA3. Thus, analysis of unit activity in CA3 will be necessary in the future. It is also interesting to extend the IG analysis such that the change of connection weights in CA1 can be detected as the influence of correlated inputs from CA3. Since there is preliminary work toward this end (Nie and Tatsuno, 2012), this extension should be possible. It is also interesting to extend the STC to include temporal lags between neurons and account for periods of partial inhibitory synchrony(Lyttle and Fellous, 2011). This way, clusters with various spatiotemporal configurations can be detected by the STC. Reactivation during sleep is a complex, dynamic phenomenon that requires more detailed investigations. Measurements of the precise dynamics of subgroups of neurons will provide a better understanding of memory consolidation mechanisms and the role of population coding in brain functions in general.

## 3 Materials and Methods

### 3.1 Behavioral paradigm and electrophysiology

Five adult male Brown Norway/Fischer 344 hybrid rats were used for 25-hour continuous recordings in a novel object task. Briefly, the recording sessions were divided into two 12-h sleep/rest sessions interrupted by 1-h behavior of the novel object exposure to yield a total recording duration of 25 h per recording session. Throughout the recording, the rats were allowed to move, eat, and sleep freely in the recording box, following their preferred sleep/wake cycle. During a 1-h exposure, four novel objects were placed at each corner of the recording box (height 42 cm, length 46.5 cm, and width 46.5 cm). The animal explored the novel objects, and the encounter with these novel objects was considered a novel experience. More details of the behavioral paradigm were described elsewhere (Ribeiro et al., 2004; Tatsuno et al., 2006).

The multi-unit recording was conducted using a microdrive with 12 independently adjustable tetrodes (Gothard et al., 1996), which was implanted above mPFC (3.2 mm anterior and 1.3 mm lateral (left) to bregma) and/or above HC (3.8 mm posterior and 2.5 mm lateral (left) to bregma). The recording was performed with Cheetah Data Acquisition Systems (Neuralynx), where the signals were band-pass filtered between 600 Hz and 6 kHz, and spike waveforms were recorded at 32 kHz. The data were off- line spike-sorted using KlustaKwik (K. D. Harris, Rutgers University), MClust (A. D. Redish, University of Minnesota), and WaveformCutter (S. Cowen, University of Arizona), and only the units with < 1% of inter-spike interval distribution falling within the 2 ms refractory period were used in the analyses. More details of electrophysiology and surgery techniques can be found elsewhere (Tatsuno et al., 2006).

### 3.2 Spike train clustering

A spike pattern is defined as a sequence of spikes that is temporally defined with an individual spike jitter 𝛿. The identification of groups relies on the choice of a bin-less distance measure between spike trains. We choose a distance obtained as the dot product of the convolutions of the discrete spike trains with a Gaussian kernel of width 2𝛿. The task of finding groups of nearby cells is reduced to finding clusters of cells in the space defined by vectors of spike train distances. We use a fuzzy-clustering method that is robust to initial conditions and converges rapidly. The main parameters of this method are 𝛿, T (time window in which spike patterns are considered), and K (number of clusters). The parameter 𝛿 can be estimated from the raw data and is typically in the 5-50 ms range (Fellous et al., 2004). Parameters T and K are related to the experimental contingencies and are determined empirically. This method has been successfully applied to multi-trial data in cortical slices in vitro, in the LGN of a visually stimulated anesthetized cat, and in area MT of an awake behaving monkey (Fellous et al., 2004).

### 3.3 Information geometry

Information Geometry (IG) investigates the geometrical structures of the parameter space of probability distributions. For example, a normal distribution 𝑝(𝑥; 𝜇, 𝜎) can be studied in the space spanned by the two parameters 𝜇 (mean) and 𝜎 (standard deviation). The study of geometrical structures of distributions of an exponential family and a mixture family reveals that the natural and expectation parameters can be decomposed orthogonally (Amari and Nagaoka, 2000; Amari, 2016). To apply the orthogonal property to multi-unit spike data, each spike train is represented by binary random variables. This allows the probability distribution of neural firing to be expanded by the log-linear model exactly. To obtain the IG measures, we first represent a state of neurons by binary variables 𝑥_*i*_ (𝑥_*i*_ = 1 for a neuron *i* firing in a short time interval and 0 otherwise). Using the log-linear model, the joint firing probability of neurons *i* and *j*, 𝑝_𝑥*i*𝑥*j*_, is expressed as

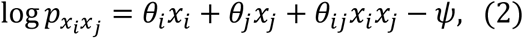

where 𝜓 is a normalization factor for ∑ 𝑝_𝑥*i*𝑥*j*_ = 1. Since 𝑥_*i*_ and 𝑥_*j*_ take on the binary values 0, 1, we have,

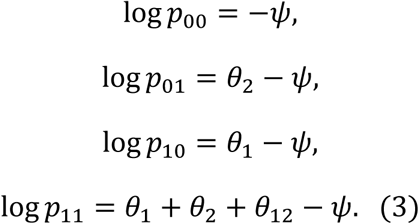

The coefficients *θ*_1_, *θ*_2_ and *θ*_12_ are then given by

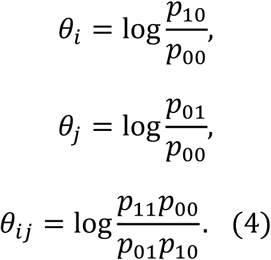

The *θ*_*i*_ and *θ*_*j*_ are called the first-order IG measures (the 1st-order IGM) and *θ*_*ij*_the second-order IG measure (the 2nd-order IGM). It has been shown that *θ*_*ij*_ and the firing rate of neurons *i* and *j*, 𝜂_*i*_ and 𝜂_*j*_, are orthogonal to each other where 𝜂_*i*_ = 𝑝_10_ + 𝑝_11_ and 𝜂_*j*_ = 𝑝_01_ + 𝑝_11_ (Amari, 2001). For IG analysis of the biophysical simulations, the probabilities 𝑝_00_, 𝑝_01_, 𝑝_10_ and 𝑝_11_ were estimated from 500 independent samples of spike trains using 20 ms bin width with Gaussian smoothing over 10 bins (Fig. 2). For the IG analysis of in vivo multi-unit data, the probabilities were estimated from spike trains within 15 min block of recording using 100 ms bin width and Gaussian smoothing over 5 bins (Fig. 4). Multiple spikes occurring in a single bin were converted to a single spike.

### 3.4 Biophysical simulation

Model neurons were single compartments containing a generic leak current, sodium and potassium currents responsible for action potential generation (Golomb and Amitai, 1997), a generic high-voltage calcium current (Reuveni et al., 1993), a first-order calcium pump (McCormick and Huguenard, 1992; Destexhe et al., 1994a) and a calcium-activated potassium current to control burst firing (Destexhe et al., 1994b). In addition, two generic background synaptic noise currents (excitatory and inhibitory) were added (Destexhe et al., 2001; Destexhe et al., 2003) to recreate in vivo conditions (Pare et al., 1998; Destexhe et al., 2001; Fellous et al., 2003). The strength of these connections is random and bounded so that no single cell can fire its postsynaptic target more than 25% of the time.

### 3.5 Explained variance

The explained variance was introduced to evaluate the similarity between the correlation matrices of the task and post-task rest epochs by taking the pre-existing correlation between the task and pre-task rest epochs into account (Kudrimoti et al., 1999). It is defined as the square of the partial correlation,

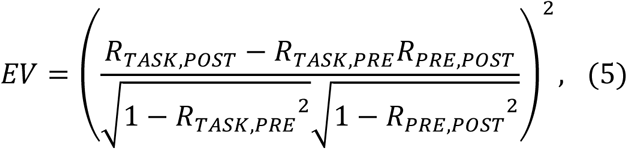

where 𝑅_𝑇𝐴𝑆𝐾,𝑃𝑅𝐸_, 𝑅_𝑇𝐴𝑆𝐾,𝑃𝑂𝑆𝑇_, and 𝑅_𝑃𝑅𝐸,𝑃𝑂𝑆𝑇_are the Pearson correlation coefficients between the correlation matrices of the task and pre-task rest epochs, between the correlation matrices of the task and post-task rest epochs, and between the correlation matrices of the pre-task rest and post-task rest epochs, respectively. The reversed EV was calculated by exchanging PRE and POST in the above EV formula (Pennartz et al., 2004). To obtain a detailed time course of the memory reactivation signal, the EV and reversed EV were computed by shifting a 15 min block of post-task and pre-task correlation matrices in a 5 min step. Significant reactivation was detected when EV exceeded the mean + 2 standard deviation of reversed EV in each block. More details of the EV analysis can be found elsewhere (Tatsuno et al., 2006).

## 5 Conflict of Interest

The authors declare that the research was conducted in the absence of any commercial or financial relationships that could be construed as a potential conflict of interest.

## 6 Author Contributions

M.T. conducted experiments, analyzed data, and wrote the manuscript. JM.F. analyzed data and wrote the manuscript.

## 7 Funding

NSF-CRCNS 1010172; RGPIN-2020-06342; AI-232403195

## Acknowledgments

The authors thank Bruce L. McNaughton for his advice and Karim Ali for his technical assistance.

## 9 Data Availability Statement

Data will be made available after publication.

## Notes

### Competing Interest Statement

The authors have declared no competing interest.

## References

Ahrens MB, Engert F (2015) Large-scale imaging in small brains. Curr Opin Neurobiol 32:78–86.

Amari S (2001) Information Geometry on Hierarchy of Probability Distributions. IEEE Transactions on Information Theory 47:1701–1711.

Amari S (2009) Measure of correlation orthogonal to change in firing rate. Neural Comput 21:960–972.

Amari S (2016) Information geometry and its application: Springer.

Amari S, Nagaoka H (2000) Methods of information geometry. New York: Oxford University Press.

Bartho P, Hirase H, Monconduit L, Zugaro M, Harris K, Buzsaki G (2004) Characterization of neocortical principal cells and interneurons by network interactions and extracellular features. J Neurophysiol.

Buzsaki G (1998) Memory consolidation during sleep: a neurophysiological perspective. J Sleep Res 7 Suppl 1:17–23.

Buzsaki G (2004) Large-scale recording of neuronal ensembles. Nat Neurosci 7:446–451.

Buzsaki G, Stark E, Berenyi A, Khodagholy D, Kipke DR, Yoon E, Wise KD (2015) Tools for probing local circuits: high-density silicon probes combined with optogenetics. Neuron 86:92–105.

Chang EY, Morris KF, Shannon R, Lindsey BG (2000) Repeated sequences of interspike intervals in baroresponsive respiratory related neuronal assemblies of the cat brain stem. J Neurophysiol 84:1136–1148.

de la Rocha J, Doiron B, Shea-Brown E, Josic K, Reyes A (2007) Correlation between neural spike trains increases with firing rate. Nature 448:802–806.

Destexhe A, Mainen ZF, Sejnowski TJ (1994a) Synthesis of models for excitable membranes, synaptic transmission and neuromodulation using a common kinetic formalism. J Comput Neurosci 1:195–230.

Destexhe A, Rudolph M, Pare D (2003) The high-conductance state of neocortical neurons in vivo. Nat Rev Neurosci 4:739–751.

Destexhe A, Contreras D, Sejnowski TJ, Steriade M (1994b) A model of spindle rhythmicity in the isolated thalamic reticular nucleus. J Neurophysiol 72:803–818.

Destexhe A, Rudolph M, Fellous JM, Sejnowski TJ (2001) Fluctuating synaptic conductances recreate in vivo-like activity in neocortical neurons. Neuroscience 107:13–24.

Fellous JM, Rudolph M, Destexhe A, Sejnowski TJ (2003) Synaptic background noise controls the input/output characteristics of single cells in an in vitro model of in vivo activity. Neuroscience 122:811–829.

Fellous JM, Tiesinga PH, Thomas PJ, Sejnowski TJ (2004) Discovering spike patterns in neuronal responses. J Neurosci 24:2989–3001.

Giocomo LM (2015) Large scale in vivo recordings to study neuronal biophysics. Curr Opin Neurobiol 32:1–7.

Giri B, Miyawaki H, Mizuseki K, Cheng S, Diba K (2019) Hippocampal Reactivation Extends for Several Hours Following Novel Experience. J Neurosci 39:866–875.

Golomb D, Amitai Y (1997) Propagating neuronal discharges in neocortical slices: computational and experimental study. J Neurophysiol 78:1199–1211.

Gothard KM, Skaggs WE, Moore KM, McNaughton BL (1996) Binding of hippocampal CA1 neural activity to multiple reference frames in a landmark-based navigation task. J Neurosci 16:823–835.

Grinvald A, Petersen CC (2015) Imaging the Dynamics of Neocortical Population Activity in Behaving and Freely Moving Mammals. Adv Exp Med Biol 859:273–296.

Hamel EJ, Grewe BF, Parker JG, Schnitzer MJ (2015) Cellular level brain imaging in behaving mammals: an engineering approach. Neuron 86:140–159.

Harris KD, Quiroga RQ, Freeman J, Smith SL (2016) Improving data quality in neuronal population recordings. Nat Neurosci 19:1165–1174.

Hebb DO (1949) The Organization of Behavior: A neuropsychological theory. New York: Wiley.

Hines ML, Carnevale NT (1997) The NEURON simulation environment. Neural Comput 9:1179-1209.

Humphries MD (2011) Spike-train communities: finding groups of similar spike trains. J Neurosci 31:2321–2336.

Kim TH, Zhang Y, Lecoq J, Jung JC, Li J, Zeng H, Niell CM, Schnitzer MJ (2016) Long-Term Optical Access to an Estimated One Million Neurons in the Live Mouse Cortex. Cell Rep 17:3385–3394.

Kudrimoti HS, Barnes CA, McNaughton BL (1999) Reactivation of hippocampal cell assemblies: effects of behavioral state, experience, and EEG dynamics. J Neurosci 19:4090–4101.

Lipa P, Tatsuno M, McNaughton BL, Fellous JM (2007) Dynamics of neural assemblies involved in memory-trace replay. Soc Neurosci Abstr 37.308:13.

Lipa P, Tatsuno M, Amari S, McNaughton BL, Fellous JM (2006) A novel analysis framework for characterizing ensemble spike patterns using spike train clustering and information geometry. Soc Neurosci Abstr 36:371:6.

Lyttle D, Fellous JM (2011) A new similarity measure for spike trains: sensitivity to bursts and periods of inhibition. J Neurosci Methods 199:296–309.

Mackevicius EL, Bahle AH, Williams AH, Gu S, Denisenko NI, Goldman MS, Fee MS (2019) Unsupervised discovery of temporal sequences in high-dimensional datasets, with applications to neuroscience. Elife 8.

McClelland JL, McNaughton BL, O’Reilly RC (1995) Why there are complementary learning systems in the hippocampus and neocortex: insights from the successes and failures of connectionist models of learning and memory. Psychol Rev 102:419–457.

McCormick DA, Huguenard JR (1992) A model of the electrophysiological properties of thalamocortical relay neurons. J Neurophysiol 68:1384–1400.

Miller EK, Wilson MA (2008) All my circuits: using multiple electrodes to understand functioning neural networks. Neuron 60:483–488.

Nakahara H, Amari S (2002) Information-geometric measure for neural spikes. Neural Comput 14:2269–2316.

Nicolelis MA, Lebedev MA (2009) Principles of neural ensemble physiology underlying the operation of brain-machine interfaces. Nat Rev Neurosci 10:530–540.

Nie Y, Tatsuno M (2012) Information-Geometric Measures for Estimation of Connection Weight Under Correlated Inputs. Neural Comput.

Nie Y, Fellous JM, Tatsuno M (2014a) Information-geometric measures estimate neural interactions during oscillatory brain states. Front Neural Circuits 8:11.

Nie Y, Fellous JM, Tatsuno M (2014b) Influence of external inputs and asymmetry of connections on information-geometric measures involving up to ten neuronal interactions. Neural Comput 26:2247–2293.

Nienborg H, Cumming B (2010) Correlations between the activity of sensory neurons and behavior: how much do they tell us about a neuron’s causality? Curr Opin Neurobiol 20:376–381.

Pare D, Shink E, Gaudreau H, Destexhe A, Lang EJ (1998) Impact of spontaneous synaptic activity on the resting properties of cat neocortical pyramidal neurons In vivo. J Neurophysiol 79:1450–1460.

Pennartz CM, Lee E, Verheul J, Lipa P, Barnes CA, McNaughton BL (2004) The ventral striatum in off-line processing: ensemble reactivation during sleep and modulation by hippocampal ripples. J Neurosci 24:6446–6456.

Pospisil DA, Bair W (2021) Accounting for Biases in the Estimation of Neuronal Signal Correlation. J Neurosci 41:5638–5651.

Prut Y, Vaadia E, Bergman H, Haalman I, Slovin H, Abeles M (1998) Spatiotemporal structure of cortical activity: properties and behavioral relevance. J Neurophysiol 79:2857–2874.

Ramirez-Amaya V, Vazdarjanova A, Mikhael D, Rosi S, Worley PF, Barnes CA (2005) Spatial exploration-induced Arc mRNA and protein expression: evidence for selective, network-specific reactivation. J Neurosci 25:1761–1768.

Rector DM, Yao X, Harper RM, George JS (2009) In Vivo Observations of Rapid Scattered Light Changes Associated with Neurophysiological Activity. In: In Vivo Optical Imaging of Brain Function (nd, Frostig RD, eds). Boca Raton (FL).

Reuveni I, Friedman A, Amitai Y, Gutnick MJ (1993) Stepwise repolarization from Ca2+ plateaus in neocortical pyramidal cells: evidence for nonhomogeneous distribution of HVA Ca2+ channels in dendrites. J Neurosci 13:4609–4621.

Ribeiro S, Gervasoni D, Soares ES, Zhou Y, Lin SC, Pantoja J, Lavine M, Nicolelis MA (2004) Long-lasting novelty-induced neuronal reverberation during slow-wave sleep in multiple forebrain areas. PLoS Biol 2:E24.

Russo E, Durstewitz D (2017) Cell assemblies at multiple time scales with arbitrary lag constellations. Elife 6.

Schwindel CD, Ali K, McNaughton BL, Tatsuno M (2014) Long-term recordings improve the detection of weak excitatory-excitatory connections in rat prefrontal cortex. J Neurosci 34:5454–5467.

Shimazaki H, Amari S, Brown EN, Grun S (2012) State-space analysis of time-varying higher-order spike correlation for multiple neural spike train data. PLoS Comput Biol 8:e1002385.

Song S, Sjostrom PJ, Reigl M, Nelson S, Chklovskii DB (2005) Highly nonrandom features of synaptic connectivity in local cortical circuits. PLoS Biol 3:e68.

Steinmetz NA, Koch C, Harris KD, Carandini M (2018) Challenges and opportunities for large-scale electrophysiology with Neuropixels probes. Curr Opin Neurobiol 50:92–100.

Stringer C, Pachitariu M (2024) Analysis methods for large-scale neuronal recordings. Science 386:eadp7429.

Stringer C, Pachitariu M, Steinmetz N, Reddy CB, Carandini M, Harris KD (2019) Spontaneous behaviors drive multidimensional, brainwide activity. Science 364:255.

Tanaka T (2000) Information geometry of mean-field approximation. Neural Comput 12:1951-1968.

Tatsuno M (2015) Analysis and modeling of coordinated multi-neuronal activity. New York: Springer.

Tatsuno M, Okada M (2004) Investigation of possible neural architectures underlying information-geometric measures. Neural Comput 16:737–765.

Tatsuno M, Lipa P, McNaughton BL (2006) Methodological considerations on the use of template matching to study long-lasting memory trace replay. J Neurosci 26:10727–10742.

Tatsuno M, Fellous JM, Amari SI (2009) Information-Geometric Measures as Robust Estimators of Connection Strengths and External Inputs. Neural Comput 21:2309–2335.

Urai AE, Doiron B, Leifer AM, Churchland AK (2022) Large-scale neural recordings call for new insights to link brain and behavior. Nat Neurosci 25:11–19.

Watanabe K, Haga T, Tatsuno M, Euston DR, Fukai T (2019) Unsupervised Detection of Cell-Assembly Sequences by Similarity-Based Clustering. Front Neuroinform 13:39.

Yuste R, Cossart R, Yaksi E (2024) Neuronal ensembles: Building blocks of neural circuits. Neuron 112:875–892.

